# An extraordinary larval-like teleost fish from the Eocene of Bolca

**DOI:** 10.1101/2024.08.19.608581

**Authors:** Donald Davesne, Giorgio Carnevale

## Abstract

“*Pegasus*” *volans* is a highly unusual fossil teleost fish from the celebrated Eocene Bolca Lagerstatte. The fossil, known on the basis of two specimens, has been historically assigned to seamoths (Pegasidae), then to oarfish and relatives (Lampriformes). We describe its enigmatic skeletal anatomy in detail, and provide a new genus name. “*Pegasus*” *volans* is an extremely elongate and slender animal, with long anal and dorsal fins and a very well-developed first dorsal-fin ray reminiscent to the vexillum of some modern teleost larvae. Most striking is its extreme ventral projection of the pelvic girdle (basipterygium), associated with an element of the pectoral girdle (a long process of the coracoid) and developed pelvic-fin rays. The strongly reduced abdominal region suggests that “*Pegasus*” *volans* had an external gut, once again reminiscent of those of certain larval teleosts. The unique character state combination displayed by “*Pegasus*” *volans* make it impossible to assign it to a specific subclade within perch-like spiny-rayed fishes (Percomorpha). Nevertheless, it offers a valuable perspective on the diversity of morphologies and ecological niches occupied by teleost fishes of the early Eocene Bolca fauna.

## INTRODUCTION

The early Eocene Bolca Lagerstatte of northern Italy famously records the earliest examples of a modern, reef-associated shallow marine fauna (Carnevale *et al*. 2014; Friedman & Carnevale 2018) with specimens showing exquisitely preserved anatomical details. The fauna includes some of the oldest known representatives of many ray-finned taxa with a major presence in shallow reefs today – such as eels (Anguilliformes), squirrelfishes (Holocentridae), cardinalfishes (Apogonidae), pipefishes and relatives (Syngnathiformes), damselfishes (Pomacentridae), wrasses (Labridae) or surgeonfishes (Acanthuridae), alongside fewer representatives of a more open-ocean ecosystem such as flyingfishes (Exocoetidae), jacks (Carangidae), or mackerels and tunas (Scombridae).

Possibly due to its palaeoecological and palaeogeographical setting within an inferred historical biodiversity hotspot (Renema *et al*. 2008; Marramà *et al*. 2016) the Bolca faunas also records morphological oddities, not found in modern ray-finned fish biodiversity. Examples include the partially asymmetrical flatfishes †*Amphistium paradoxum* and †*Heteronectes chaneti* (Friedman 2008), the elongate relative of modern dories †*Bajaichthys elegans* (Davesne *et al*. 2017), the shallow-water barracudina †*Holosteus* (Marramà & Carnevale 2017), and the long-snouted †Rhamphosidae, heavily spined relatives of the modern flying gurnards (Calzoni *et al*. 2023).

In this paper, we describe in detail the anatomy of an extraordinary and enigmatic fossil from Bolca which shows features commonly observed today in teleost larvae. Although known since the 18^th^ century, the taxon has remained enigmatic ever since. For instance, Blot (1980) refrained from providing a systematic attribution in his complete review of the Bolca ichthyofauna. More recently, some authors tentatively referred the taxon to Lampriformes (Bannikov 2014), more specifically to the elongate Taeniosomi that include oarfishes, ribbonfishes and crestfishes (Carnevale *et al*. 2014; Carnevale & Bannikov 2018), but without anatomical arguments to support such attribution.

## MATERIAL AND METHODS

Only two specimens are known. The holotype, MNHN.F.BOL65/BOL66, in part and counterpart (Fig. 1), is housed at the Muséum national d’Histoire naturelle (MNHN), Paris, France. The other specimen, MCSNV T.293/T.294, also in part and counterpart (Figs. 2-5) belongs to the Baja collection housed at the Museo Civico di Storia Naturale (MCSNV), Verona, Italy. Both specimens show the anterior portion of the body, including the head, but the posterior end is not preserved. They are remarkably similar in morphology and are within the same size range (approximately 6.6 mm head length for MNHN.F.BOL65/BOL66 and 5.7 mm head length for MCSNV T.293/T.294).

**Figure 1.**
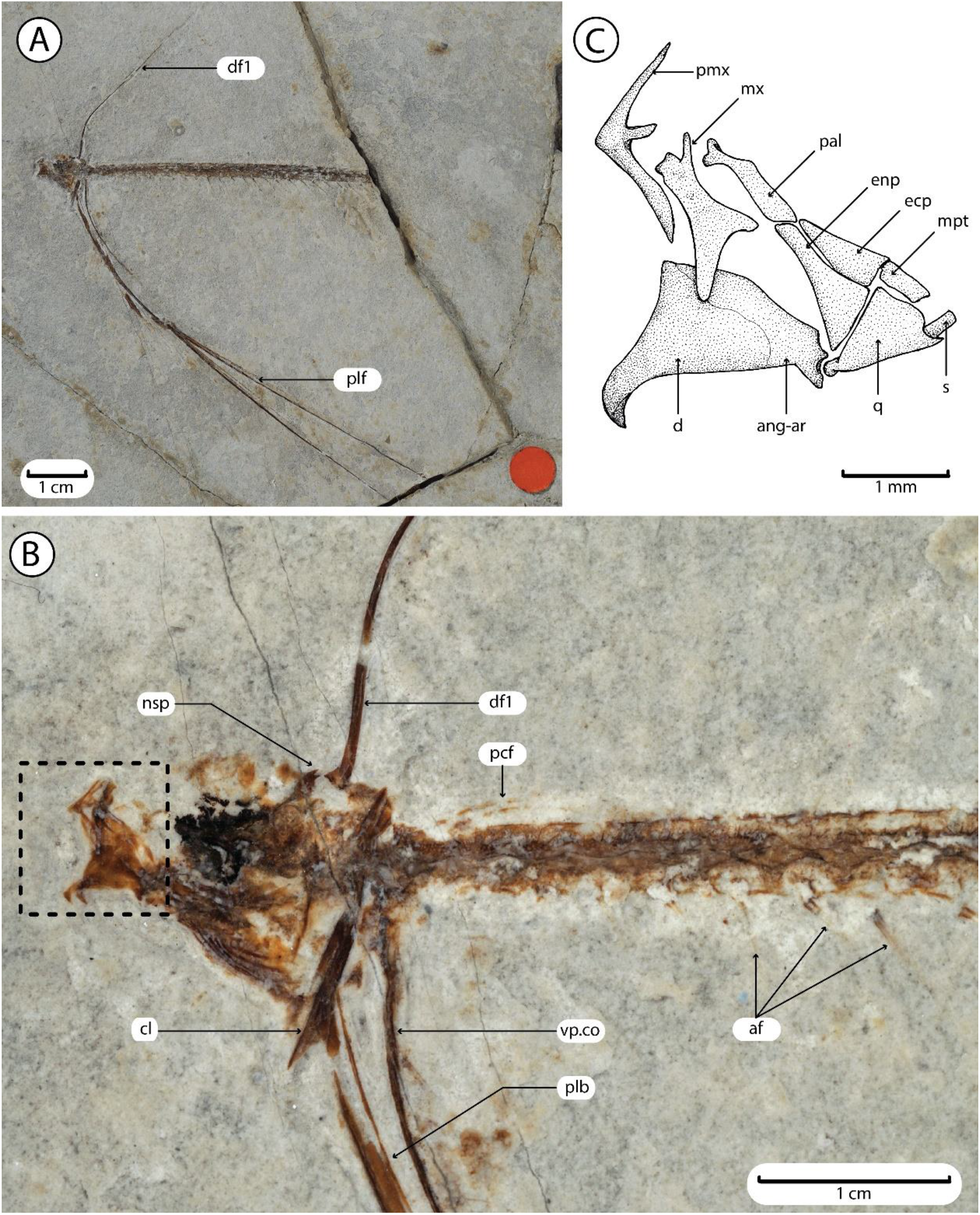
“*Pegasus*” *volans*, holotype MNHN.F.BOL65. **A:** Photograph of the entire specimen. **B:** Detail of the cephalic and pectoral regions, moistened with alcohol. **C:** Interpretative drawing of the suspensorium and jaws. Counterpart MNHN.F.BOL66 is poorly preserved and not figured here. Photographs by P. Loubry (MNHN). **Legend:** af, anal-fin rays; ang-ar, anguloarticular; cl, cleithrum; d, dentary; df1, dorsal-fin ray 1 (vexillum-like); ecp, ectopterygoid; enp, endopterygoid; mpt, metapterygoid; mx, maxilla; pal, palatine; nsp, bifid spine at the postero-dorsal corner of the neurocranium; pcf, pectoral-fin rays; plb, pelvic bone (basipterygium); plf, pelvic-fin rays; pmx, premaxilla; q, quadrate; s, symplectic; vp.co, ventral process of the coracoid.

**Figure 2.**
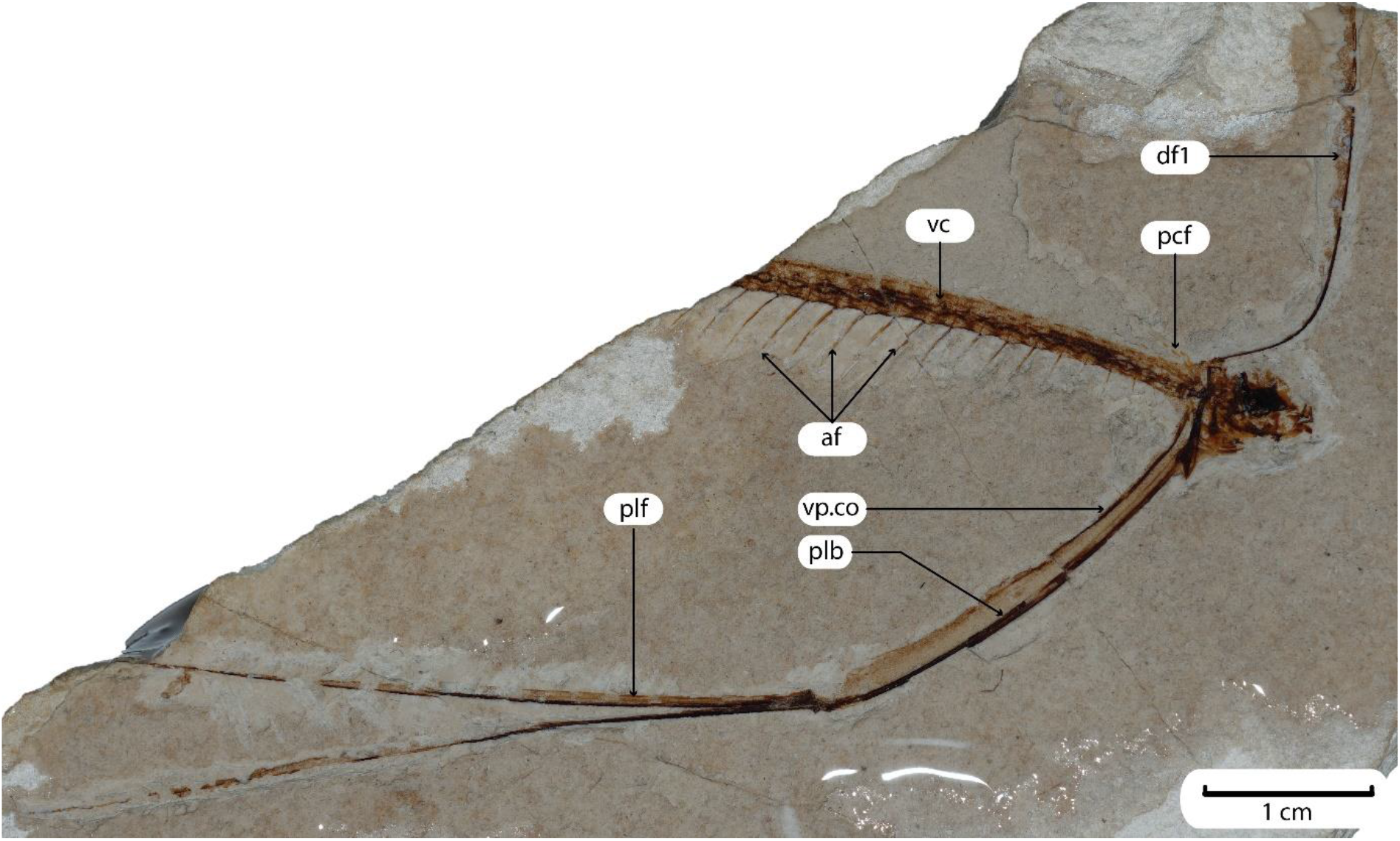
“*Pegasus*” *volans*, MCSNV T.293. Photograph of the entire specimen. Photograph by D. Davesne. **Legend:** af, anal-fin rays; df1, dorsal-fin ray 1 (vexillum-like); pcf, pectoral-fin rays; plb, pelvic bone (basipterygium); plf, pelvic-fin rays; vc, vertebral column; vp.co, ventral process of the coracoid.

The fossil taxon was initially described by Volta (1796). In the original cursory mention, Volta named it “*Pegasus Volans*”, which a junior synonym of *Pegasus volitans* (Fricke *et al*. 2024), an extant seamoth (Pegasidae, Syngnathiformes) from the Indo-Pacific region. There is very little in common morphologically between *Pegasus* and the fossils studied herein: the former is a benthic, dorsoventrally flattened animal with an elongate rostrum, conspicuous bony plates and well-developed pectoral fins. On the contrary, the fossil described by Volta is a laterally flattened animal with no rostrum or bony plates. Despite these obvious differences, the name *Pegasus volans* has been used in the literature since (Agassiz 1833; Blot 1980) – even though recent authors have acknowledged the need for a new genus and referred it to as “*Pegasus*” *volans* (Bannikov 2014; Carnevale *et al*. 2014; Brignon 2019).

For this study, fossils were re-examined using a Leica M80 stereomicroscope equipped with a camera lucida drawing arm. To enhance details, specimens were moistened with alcohol. Photographs were taken under UV light to reveal potentially hidden structures.

## SYSTEMATIC PALAEONTOLOGY

Teleostei Müller, 1845

Acanthomorpha Rosen, 1973

Percomorpha Cope, 1871

Percomorpha *incertae sedis*

Genus [REDACTED] gen. nov.

### Diagnosis

Body narrow and slender. Premaxilla with a long and pointed ascending process and a well-developed and distally spatulate articular process. Maxilla large and hourglass shaped. Dentary with a prominent symphyseal spine. Oral jaw teeth tiny and conical. At least six sabre-like branchiostegal rays. Three abdominal vertebrae. First dorsal-fin ray vexillum-like, extremely elongate, flexible, unsegmented and unbranched. Dorsal and anal-fin rays unbranched and unsegmented. Anal-fin rays longer than their dorsal-fin counterparts and gradually increasing in length posteriorly. Dorsal- and anal-fin pterygiophores horizontally-oriented, hourglass-shaped. One-to-one relationship between median-fin pterygiophores and vertebrae. Cleithrum massive, straight and almost vertically-oriented. Coracoid small, bearing a remarkably elongate ventral process almost parallel to the basipterygium. Basipterygium hypertrophied protruding ventrally from the body wall for a length equal to that of the anterior 20 vertebrae. Pelvic fin with a greatly elongate, robust and distally flexible spinous ray plus two delicate unbranched and unsegmented rays. Entire body covered with tiny dermal spinules characterized by a stellate basal plate supporting an upright spine.

### Type and only species

[REDACTED] *volans* comb. nov.

**[REDACTED] *volans* comb. nov**.

1796 *Pegasus volans* Volta p. 174 pl. 42 fig. 2

1833 *Pegasus volans* Agassiz p. 35

1980 *Pegasus volans* Blot p. 384

2014 “*Pegasus*” *volans* Bannikov p. 26

2014 “*Pegasus*” *volans* Carnevale *et al*. p. 42

2018 “*Pegasus*” *volans* Carnevale & Bannikov p. 175

2019 “*Pegasus*” *volans* Brignon p. 43

### Taxonomic note

In order to guarantee priority of the new genus name for the final publication, we will continue using the obsolete name “*Pegasus*” *volans* in the remainder of the present preprint.

### Diagnosis

As for the genus.

### Holotype

MNHN.F.BOL65/BOL66, a partially complete articulated skeleton lacking the posterior portion of the body (Fig. 1), in part and counterpart.

### Referred specimens

MCSNV T.293/T.294, a partially complete articulated skeleton, lacking the posterior portion of the body (Figs. 2-5), in part and counterpart.

## DESCRIPTION

Both the available specimens are incomplete and lack the posterior part of the body, the caudal skeleton is therefore unknown. The body is extremely narrow and slender.

### Skull

The skull roof is notably inflated, apparently due to the presence of a frontal crest, probably bilateral (Fig. 3). The frontal crest has a gently rounded outer profile. It is difficult to determine whether an occipital crest and spine were present or not. The holotype has a bifid spine at the postero-dorsal corner of the neurocranium (Fig. 1). This spine is in close proximity to the base of the greatly elongate vexillum-like dorsal-fin ray.

**Figure 3.**
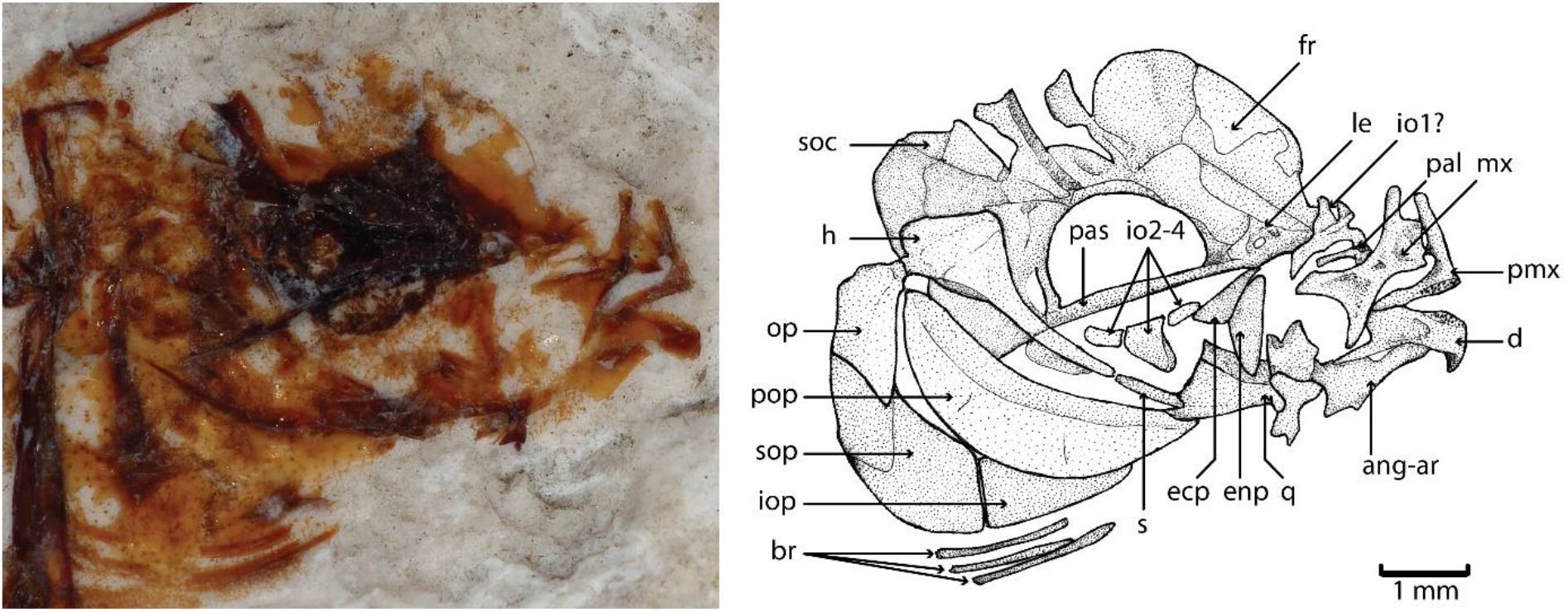
“*Pegasus*” *volans*, MCSNV T.293. Detail of the skull, and interpretative drawing. **Legend:** ang-ar, anguloarticular; br, branchiostegal rays; d, dentary; ecp, ectopterygoid; enp, endopterygoid; fr, frontal; h, hyomandibula; io, infraorbitals; iop, interopercle; le, lateral ethmoid; mx, maxilla; op, opercle; pal, palatine; pas, parasphenoid; pmx, premaxilla; pop, preopercle; q, quadrate; s, symplectic; soc, supraoccipital; sop, subopercle.

The configuration of the ethmoid region is only partially recognizable (Fig. 3), showing what appears to be the lateral ethmoid, which is in close association with the anterior articular end of the palatine.

The parasphenoid is a stout median bone that forms most of the basicranium, running in the lower half of the orbit. The basioccipital is visible only in MCSNV T.294 (Fig. 5); it is very thick and bears a dorsal ascending process that articulates with the exoccipital. The bones of the otic region of the neurocranium cannot be properly recognized due to inadequate preservation.

The eyeball is marked in both specimens by a circular, thin and dark pigmented film (Figs. 1, 3-4).

What appear to be the second, third and fourth infraorbital bones can be recognized in MCSNV T.293 (Fig. 3). These bones are feebly observable ventral to the pigmented orbit, due to their limited ossification. While the second and third elements are almost tubular, the third infraorbital is ventrally expanded.

### Jaws

The gape of the mouth is moderately developed (Figs. 1, 3). The premaxilla has a long and pointed ascending process, approximately equal in length with the alveolar process (Figs. 1, 3). There is a well-developed and distally spatulate articular process forming a deep notch with the posterior margin of the ascending process, which possibly accommodated a large rostral cartilage in origin. The premaxilla bears very small conical teeth arranged in multiple rows, which are recognizable only in MCSNV T.293 (Fig. 3).

The maxilla is toothless, large, apparently excluded from the border of the mouth, almost hourglass-shaped, and bearing a thick antero-dorsal process and concave dorsal, ventral and posterior margins (Figs. 1, 3). There is no evidence of a supramaxilla.

The mandible is almost triangular in outline (Figs. 1, 3). The dentary bears a prominent pointed and postero-ventrally directed symphysial spine. Numerous tiny conical teeth, similar to those of the premaxilla, can be observed along the alveolar process of MCSNV T.293. The angulo-articular is irregular in outline. There is a thick spine emerging from the postero-ventral corner of the mandible (Fig. 1).

### Suspensorium

Overall, the suspensorium is inclined forward (Fig. 1). The quadrate is almost triangular in shape and bears a thick articular condyle. There is a small postero-dorsal notch to accommodate the rod-like symplectic. The ectopterygoid and endopterygoid are partially preserved in both specimens, while what appears to be a quadrangular metapterygoid can be recognized in the holotype (Fig. 1). There is a prominent palatine, bearing a thick and ventrally directed anterior process. The hyomandibula has a long and slender ventral shaft and a triangular dorsal portion, with two articular heads (Fig. 3).

### Opercular region

The opercular bones are thin and only partially recognisable in the holotype (Fig. 1). The two large arms of the preopercle form an obtuse angle; the anterior margin of the bone is thickened, while the posterior margin is distinctly rounded. The opercle appears to be subrectangular in outline, with an oblique ventral margin. Interopercle and subopercle are laminar, recognizable in MCSNV T.293 (Fig. 3).

### Hyoid apparatus

The hyoid bar is compact and shows a ventral concavity at its midlength (Fig. 1). Six sabre-like branchiostegal rays appear to be present in the holotype, although their exact number is unclear due to inadequate preservation (Figs. 1, 3).

### Vertebral column

There are three abdominal vertebrae (Figs. 1-2, 4-5). Twenty-four caudal vertebrae are preserved in the holotype and 18 in MCSNV T.293 (Figs. 1-2). In both specimens the posteriormost portion of the body is not preserved, preventing the count of the original vertebral number.

The abdominal and first two caudal vertebrae are compact, shorter than the other vertebrae, and gradually increasing in size posteriorly (Figs. 4-5). The subsequent vertebrae are almost equal in size. The vertebral centra are hourglass-shaped and significantly longer than high (Figs. 4-5). The neural and haemal arches are weakly ossified, antero-posteriorly expanded, extensively ornamented with circular pits and irregular in outline. Short dorsal and ventral pre- and post-zygapophyses can be recognized. The very short neural and haemal spines are thin, pointed and vertically oriented. What appears to be a short pair of ribs can be recognized ventrally to the second abdominal vertebra in MCSNV T.293 (Fig. 4). There is no evidence of intermuscular bones.

**Figure 4.**
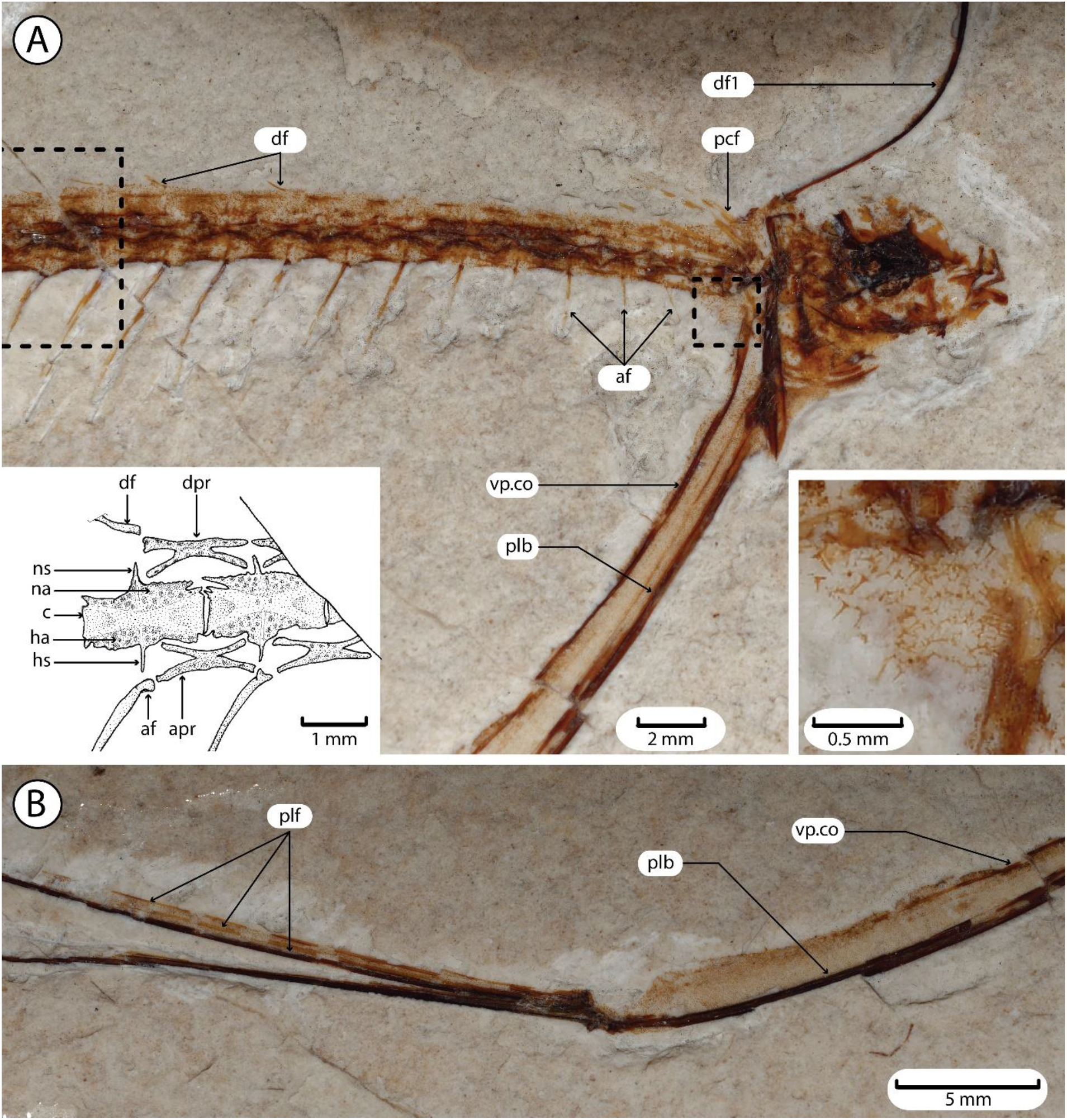
“*Pegasus*” *volans*, MCSNV T.293. Details of the postcranium. **A:** Skull, pectoral region and vertebral column. Left inset: Interpretative drawing of vertebrae 17-18 and the adjacent dorsal and anal fins and supports. Right inset: Detail of the squamation at the level of the abdomen. **B:** Distal end of the pelvic bone, and proximal end of the pelvic-fin rays. **Legend:** af, anal-fin ray; apr, anal-fin pterygiophore; c, vertebral centrum; df, dorsal-fin ray; dpr, dorsal-fin pterygiophore; ha, haemal arch; hs, haemal spine; na, neural arch; ns, neural spine; plb, pelvic bone (basipterygium); plf, pelvic-fin rays; vp.co, ventral process of the coracoid.

### Median fins and supports

There is no evidence of supraneurals. The first element of the dorsal fin is closely associated to the occipital region of the neurocranium (Figs. 1-2, 5). It is an extremely elongate, flexible, unpaired, unsegmented and unbranched stout ray, which is supported by a short but relatively thick, almost horizontal pterygiophore. In the holotype, this elongate and flexible ray appears fairly complete and tapers at its distal extremity (Fig. 1). In MCSNV T.293, it is associated with remains of a band of soft tissue that probably formed a flap along its posterior margin (Fig. 2). Numerous melanophores can be recognized along this band, which was probably pigmented in life. The rest of the dorsal fin is separated from the elongate ray by a wide gap, so that the second dorsal-fin ray inserts just above the neural spine of the fourth caudal vertebra (Fig. 4). The dorsal-fin rays are short, thin, and unsegmented, supported by horizontally-oriented pterygiophores, which are hourglass-shaped, bifurcated anteriorly and posteriorly, possibly representing fused proximal and middle radials (Fig. 4).

The first anal-fin ray inserts on the first caudal vertebra (Fig. 4). Anal-fin rays are unsegmented, notably longer than their dorsal-fin counterparts and gradually increase in length posteriorly in the series. The anal-fin pterygiophores have a similar morphology to their dorsal-fin counterparts, but they are more robustly ossified (Fig. 4).Dorsal- and anal-fin pterygiophores bear a one-to-one relationship with the vertebrae (Fig. 4).

### Pectoral girdle and fin

The posttemporal is anteriorly bifurcated and articulates ventrally with a thick and almost straight supracleithrum (Fig. 5). The cleithrum is massive, straight and almost vertical with a pointed ventral end and a moderately developed postero-dorsal flange (Figs. 1, 5). The postcleithra and the scapula are not clearly recognizable. The small coracoid articulates with the posterior margin of the cleithrum just below the vertebral column. It bears a massive and remarkably long ventral process, which runs roughly parallel to the basipterygium extending ventrally for about two thirds of the length of the basipterygium (Figs. 1--2, 5).There are five elongate pectoral-fin rays, which extend posteriorly almost to the sixth caudal vertebra (Figs. 1, 4-5). The pectoral-fin radials are not clearly visible, but the pectoral fin inserts high on the flanks, just above the level of the vertebral column.

**Figure 5.**
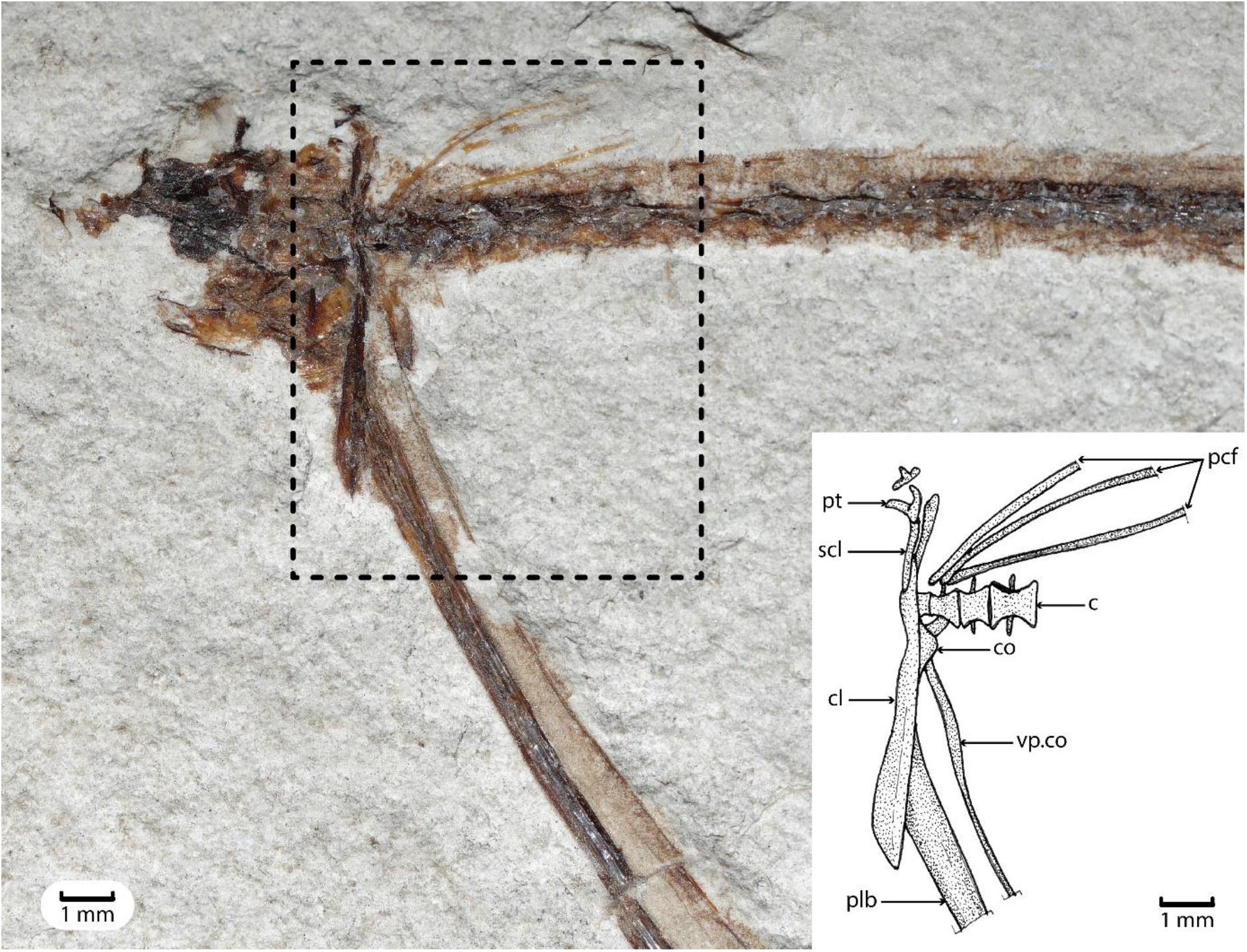
“*Pegasus*” *volans*, MCSNV T.294. Photograph of the entire specimen, and interpretative drawing of the pectoral girdle region. Photograph by D. Davesne. **Legend:** c, centra of vertebrae 1-4; cl, cleithrum; co, coracoid; pcf, pectoral-fin rays; plb, pelvic bone (basipterygium); pt, posttemporal; scl, supracleithrum; vp.co, ventral process of the coracoid.

### Pelvic girdle and fin

The basipterygium is remarkably developed and strongly ossified, projecting ventrally from the body wall, reaching a length comparable to that of the anterior 20 vertebrae (Figs. 1-2, 4-5). It gently curves ventrally and posteriorly, and seems to bear ventral and dorsal carinae at its distal extremity, giving it an oar-like appearance. Its proximal end is expanded and articulates with the cleithrum at approximately three quarters of its vertical length (Figs. 1, 4-5). The pelvic fin contains a greatly elongate, robust and distally flexible stout ray plus two unbranched and unsegmented delicate rays (Figs. 1-2, 4). The pelvic-fin rays are associated with a pigmented band of soft tissue similar to that described above for the stout first dorsal-fin ray (Fig. 4).

### Squamation and soft tissues

The whole body, including the head, median and pelvic fins, and the abdominal region (Fig. 4), is covered with tiny dermal spinules. Each spinule is characterized by a stellate basal plate supporting an upright spine. These spinules are often associated in densely packed clusters, especially around the abdominal area (Fig. 4). Examination of the holotype under UV light reveals the presence of pigmented blotches below the abdominal vertebrae and behind the (ventral process of the) coracoid-basipterygium complex.

## DISCUSSION

### Larval-like features

As described above, the ribbon-like body of “*Pegasus*” *volans* is accentuated by a stout, flexible, unsegmented and unbranched dorsal-fin ray, and by remarkably developed pelvic fins and girdle, the latter running parallel to, and associated with an exceptionally elongate ossified ventral process of the coracoid.

The elongate and flexible first dorsal-fin ray, inserting on a thick pterygiophore and probably associated with a flap of skin in life, is in some ways reminiscent of the developed first dorsal-fin rays found in various teleost larvae such as some ribbonfishes and crestfishes (Lampriformes, Trachipteridae, Lophotidae), pearlfishes (Ophidiiformes, Carapidae), cusk-eels (Ophidiiformes, Ophidiidae), flatfishes (Pleuronectoidei) or basslets (Liopropomatidae)(Amaoka 1972; Gordon *et al*. 1984; Govoni *et al*. 1984; Olney 1984; Nonaka *et al*. 2021; Girard *et al*. 2023). Most spectacularly, in larval pearlfishes this so-called “vexillum” is extremely long and ornamented with fleshy extensions (Olney & Markle 1979; Markle & Olney 1980, 1990; Gordon *et al*. 1984; Govoni *et al*. 1984).

In both specimens of “*Pegasus*” *volans* there are only three abdominal vertebrae and the body is very narrow and slender, suggesting that the abdominal cavity was extremely reduced in volume (Fig. 4). In addition, the elongate coracoid-basipterygium complex provided a ventral protrusion of the body wall. To reconcile these observations, we propose that viscera may have protruded from the body and along this ventral structure (Fig. 6), like in the exterilium (external) guts found in a diverse set of larval teleosts, including: conger eels (Anguilliformes, Congridae; Miller 2009, 2023), lanternfishes (Myctophiformes, Myctophidae; Moser *et al*. 1984), dragonfishes (Stomiiformes, Stomiidae; Kawaguchi & Moser 1984), flatfishes (Pleuronectoidei, Bothidae, Cynoglossidae, Samaridae; Amaoka 1972; Fraser & Smith 1974), pearlfishes (Ophidiiformes, Carapidae; Markle & Olney 1990) and cusk-eels (Ophidiiformes, Ophidiidae; Gordon *et al*. 1984; Fahay & Hare 2003; Fahay & Nielsen 2003; Okiyama & Yamaguchi 2004; Fukui & Kuroda 2007; Suntsov 2007; Nonaka *et al*. 2021; Girard *et al*. 2023). Remarkably, at least in cusk-eels (Ophidiidae) the exterilium gut is supported by a very long descending process of the coracoid (Okiyama & Yamaguchi 2004; Suntsov 2007; Girard *et al*. 2023), similar to that observed in “*Pegasus*” *volans*.

**Figure 6.**
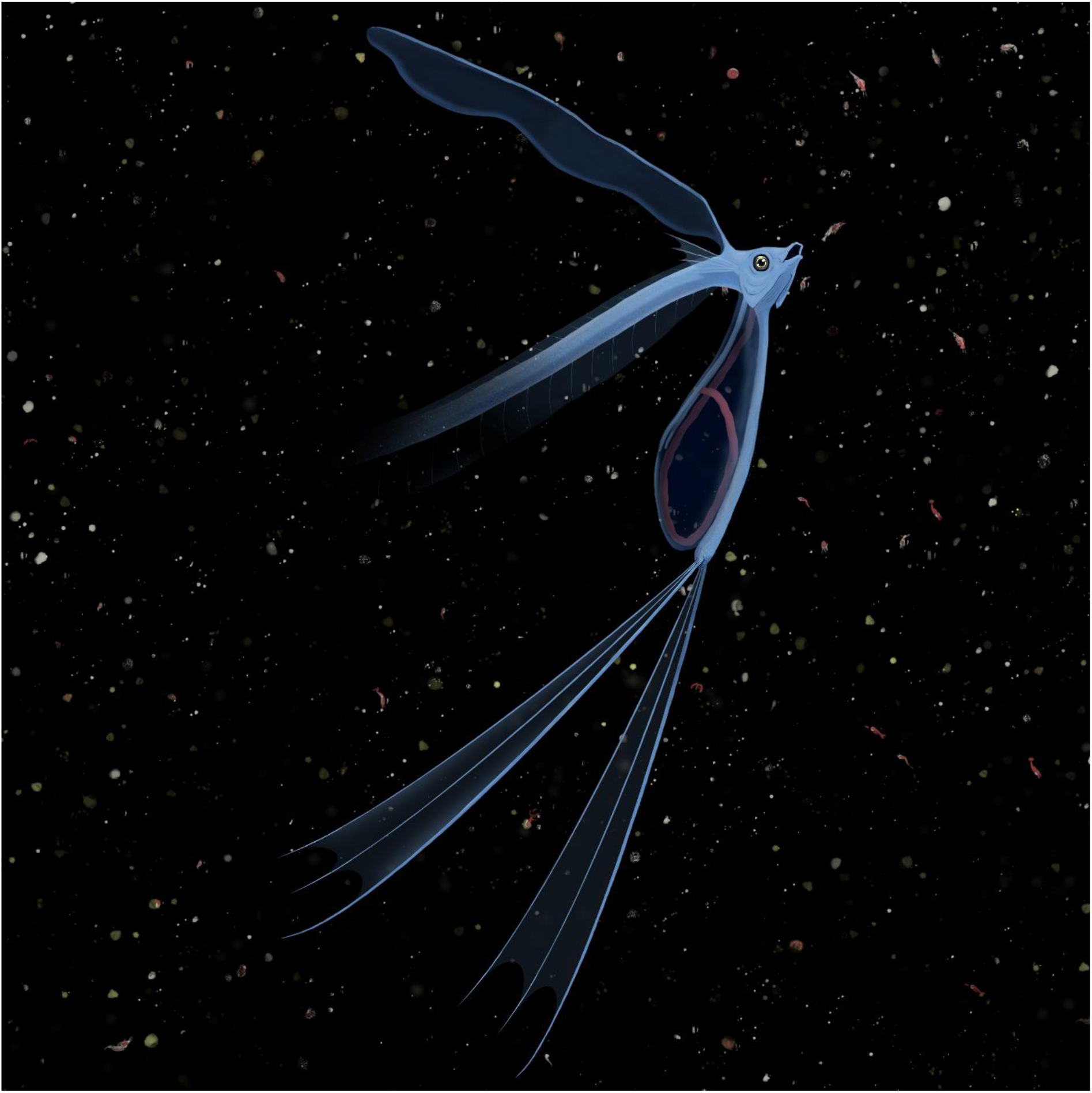
Artistic reconstruction of a living “*Pegasus*” *volans* feeding on bioluminescent plankton at night. Notice the developed vexillum-like first dorsal-fin ray, the elongate pelvic fins, and the putative external gut (exterilium) supported by the elongate ventral processes of the coracoid. Digital painting by Margaux Boetsch.

Based on these traits, which are found today exclusively in teleost larvae, “*Pegasus*” *volans* appears to have had a larval-like appearance (Fig. 6). Although teleost larvae are rare in the fossil record, a few are known from Bolca and in other Palaeogene localities (Blot & Tyler 1990; Carnevale & Bannikov 2021). In addition, these larval traits (namely the vexillum and probable exterilium) are characteristic of open-ocean pelagic larvae, and in certain cases of deep-sea fishes (e.g., cusk-eels, dragonfishes). Given that the Bolca localities record shallow marine biotopes, the occurrence of a taxon reminiscent of pelagic modern relatives is particularly striking (Fig. 6).

Although its morphology is larval-like, both the specimens of “*Pegasus*” *volans* have a similar, relatively large body size (respectively 56 and 33 mm in length, with a missing posterior extremity of unknown length). Moreover, the skeleton appears to be entirely ossified while teleost larvae usually lack ossification in at least parts of the skeleton. Therefore, we consider it dubious that the known “*Pegasus*” *volans* specimens were larvae – they may instead have been adults of a highly paedomorphic taxon retaining traits exclusive to larvae of modern teleosts.

### Functional implications

The very unusual morphology of “*Pegasus*” *volans* means that any interpretation of its ecology is highly hypothetical. Nevertheless, comparisons can be made with extant teleosts, larvae or adults, with similar morphological adaptations.

Filamentous appendages made of elongate fin rays (such as the vexillum), or even skin flaps associated with the exterilium gut, often adorn the pelagic larvae of teleost fishes, especially acanthomorphs (Moser 1981; Kendall *et al*. 1984; Nonaka *et al*. 2021). Much conjecture has surrounded the possible function of these appendages and several hypotheses have been formulated, including camouflage or predator deception (e.g., Batesian mimicry of gelatinous macroplankton, or possibly bird feathers), sensory perception and prey attraction (Moser 1981; Govoni *et al*. 1984; Kendall *et al*. 1984; Suntsov 2007; Greer *et al*. 2016; Miller 2023). Given the evidence that a skin flap was present alongside the elongate vexillum-like ray in “*Pegasus*” *volans* (Figs. 2, 6), it is possible that its developed fin rays also had similar functions.

Outside of this larval-like morphology, extreme elongation of the pelvic basipterygium is a rare feature amongst modern teleosts. A striking example can be found in the three-tooth pufferfish *Triodon macropterus* (Tetraodontiformes, Triodontidae), in which the basipterygium is mobile and supports a “pelvic fan” that can be flared and inflated to deter predators (Bemis *et al*. 2023). However, a similar function can be ruled out for “*Pegasus*” *volans*: there is no trace of a “fan” and soft-tissue preservation suggests on the contrary that skin was only covering the coracoid-basipterygium complex and its immediate vicinity (Fig. 4). Moreover, the ventral projection of the robust cleithrum and of the elongate ventral process of the coracoid that runs parallel to the pelvic basipterygium suggest that the latter was not very mobile – unlike in *Triodon* where it can perform 80° rotations (Bemis *et al*. 2023). In the absence of a clear modern analogue, the function of this elongate pelvic complex therefore remains unknown.

### Phylogenetic affinities

Descriptive analysis of the skeletal anatomy of “*Pegasus*” *volans* shows a highly unusual combination of morphological features, making it challenging to determine its phylogenetic affinities within teleosts. However, it is evident that this peculiar Eocene genus is not closely related to the modern seamoth *Pegasus* and, more generally, to the syngnathiform family Pegasidae (Santaquiteria *et al*. 2021). Pegasidae contains two extant genera (*Eurypegasus* and *Pegasus*) of benthic fishes characterized by a unique body plan, consisting of a depressed body completely encased in bony plates, a small head bearing a well-developed nasal rostrum protruding anteriorly far beyond the mouth, and large, wing-like and horizontally oriented pectoral fins (Pietsch 1978; Palsson & Pietsch 1989). None of these features can be recognized in “*Pegasus*” *volans*, unquestionably showing the lack of a close affinity with Pegasidae.

Despite the absence of true spines in the dorsal, anal and pelvic fins (Rosen 1973; Johnson & Patterson 1993), “*Pegasus*” *volans* should be regarded as a spiny-rayed teleost (Acanthomorpha) based on the very anterior position of its dorsal fin and on the contact between its pelvic and pectoral girdles (Stiassny & Moore 1992; Johnson & Patterson 1993; Parenti & Song 1996; Davesne *et al*. 2016). In addition, the overall configuration of the jaw bones, with a large premaxilla bearing well-developed ascending and articular process and a large toothless maxilla excluded from the relatively short gape of the mouth also support the attribution of “*Pegasus*” *volans* to the acanthomoph clade (Schaeffer & Rosen 1961; Patterson 1964; Rosen & Patterson 1969; Johnson & Patterson 1993; Davesne *et al*. 2016). Acanthomorphs underwent a massive evolutionary radiation in the aftermath of the Cretaceous-Palaeogene extinction event, and while their Paleocene fossil record is still poorly known and understood (Friedman 2010; Alvarado-Ortega *et al*. 2015; Argyriou & Davesne 2021; El-Sayed *et al*. 2021; Friedman *et al*. 2023; Schwarzhans *et al*. 2024) it is clear that most of the extant acanthomorph lineages were present by the early Eocene (Friedman & Carnevale 2018; Friedman *et al*. 2023; Schwarzhans *et al*. 2024).

Due to its general resemblance to certain larval and juvenile taeniosome lampriforms such as ribbonfishes, crestfishes and oarfishes (Olney 1984; Oka *et al*. 2020), Bannikov (2014) and Carnevale *et al*. (2014) tentatively referred “*Pegasus*” *volans* to the Taeniosomi. Lampriformes are known in Bolca with two genera of Veliferidae (Bannikov 1990; Carnevale & Bannikov 2018), and the earliest known fossil taeniosomes hail from slightly younger localities of Iran and the Caucasus (Daniltshenko 1980; Davesne 2017). While “*Pegasus*” *volans* exhibits certain features that also occur in lampriforms (well-developed ascending process of the premaxilla, supramaxillae absent, less than seven branchiostegal rays), it lacks the majority of the lampriform synapomorphies (Olney *et al*. 1993; Davesne *et al*. 2014, 2016, 2023; Delbarre *et al*. 2016), including: the absence of the anterior palatine process, the presence of frontal vault to accommodate the premaxilla and rostral cartilage, and the mesethmoid posterior to the lateral ethmoids. In addition, modern taeniosome lampriforms have a reduced or absent anal fin (Olney 1984; Olney *et al*. 1993), in contrast to the very developed anal fin of “*Pegasus*” *volans*. Therefore, based on the available evidence, the attribution of “*Pegasus*” *volans* to Lampriformes can be ruled out.

Another taxon from Bolca with superficial similarities to “*Pegasus*” *volans* is †*Bajaichthys elegans*, formerly interpreted as a lampriform and now referred to the Zeiformes (Sorbini & Bottura 1988; Davesne *et al*. 2017). Similarities include an elongate body with a long anal fin base, an anteriorly-inserted dorsal fin, developed pelvic-fin rays and unbranched dorsal and anal-fin rays. However, †*Bajaichthys* and zeiforms in general show dorsal, anal and pelvic-fin spines which are entirely absent in “*Pegasus*” *volans*, and no known zeiform shows the extreme vertical expansion of the coracoid and of the basipterygium observed in “*Pegasus*” *volans*. Other differences with †*Bajaichthys* include the dorsal fin running along the whole body, the absence of supraneurals, and much less developed jaws and suspensorium.

Among the peculiar diagnostic features of “*Pegasus*” *volans*, the stout and flexible vexillum-like first dorsal-fin ray and the possibly occurrence of an exterilium gut are shared by certain ophidiiform larvae (Fraser & Smith 1974; Olney & Markle 1979; Gordon *et al*. 1984; Govoni *et al*. 1984; Markle & Olney 1990; Girard *et al*. 2023). In particular, the ventral projection of the coracoid supporting the exterilium gut is seemingly unique to ophidiiform larvae in modern teleosts (Okiyama & Yamaguchi 2004; Suntsov 2007; Girard *et al*. 2023), in which it is strongly reminiscent of the structure observed in “*Pegasus*” *volans*. Ophidiiformes (cusk-eels and pearlfishes) are known in the fossil record from the Late Cretaceous onward (Carnevale & Johnson 2015), and are represented in Bolca by “*Ophidium*” *voltianum*, a potential ophidiid that is in dire need of redescription (Bannikov 2014; Carnevale *et al*. 2014). The few osteological synapomorphies that have been identified for Ophidiiformes include the lack of ossified supraneurals (Patterson & Rosen 1989), a supraoccipital that is excluded from the posterior margin of the skull by dorsally enlarged exoccipitals (Carnevale & Johnson 2015), and a reduced second infraorbital overlapping the dorsal margin of the first infraorbital (Ohashi 2018). While supraneurals are also absent in “*Pegasus*” *volans*, the other two characters cannot be clearly observed due to inadequate preservation of the cranial region in both specimens. In addition, modern ophidiiforms tend to have reduced or entirely absent pelvic fins containing a maximum of one or two rays in both larvae and adults (Gordon *et al*. 1984), while “*Pegasus*” *volans* shows strongly developed pelvic-fin rays. An attribution to Ophidiiformes, while tempting due to close similarities with larval representatives of the clade, is therefore difficult to ascertain based on available data.

The dorsal- and anal-fin pterygiophores of “*Pegasus*” *volans* likely consist of fused proximal and middle radials and appear to bear a one-to-one relationship with the adjacent vertebrae, showing a condition exclusive to certain percomorph acanthomorphs, including blennies (Blennioidei), roundheads and eel-blennies (Plesiopidae), bandfishes (Cepolidae), and eelpouts (Zoarcidae) amongst others (Gosline 1968; Smith-Vaniz & Johnson 1990; Springer 1993; Mooi & Gill 2004). Conversely, this one-to-one relationship appears to not be found in modern ophidiiforms (Gordon *et al*. 1984). These groups of mostly benthic percomorphs commonly exhibit unsegmented median-fin rays, similar to those observed in “*Pegasus*” *volans* (Gosline 1968). Finally, the extensive cover of tiny upright dermal spinules emerging from stellate basal plates characteristic of “*Pegasus*” *volans* may provide a further indication of its percomorph affinities, since modified plates and spinules are most commonly found in certain representatives of the clade (Roberts 1993).

On the basis of the available evidence, the precise phylogenetic position of “*Pegasus*” *volans* remains uncertain. However, its combination of character states support an attribution to Percomorpha, the major Cenozoic radiation of acanthomorphs that taxonomically dominate the Bolca fauna (Friedman & Carnevale 2018). While its ecology cannot be inferred due to its unique morphology, similarities with modern pelagic larvae suggest that this animal may have been associated with a more open-ocean environment, which could explain its rarity amongst fossil teleosts from Bolca.

## ACKNOWLEDGEMENTS

We would like to thank the curators that granted access to the specimens of study: Gaël Clément, Alan Pradel (MNHN), and Roberto Zorzin (MCSNV). Alexandre Bannikov (Borissiak Paleontological Institute), Matt Friedman (University of Michigan), Chase Brownstein (Yale University) and James Andrews (University of Michigan) are thanked for their insights on the fossil taxon studied here. Finally, we would like to thank Philippe Loubry (MNHN) for providing photographs of the MNHN specimens, and Margaux Boetsch for the wonderful artistic reconstruction. The research of GC was supported by a SYNTHESIS grant and by grants (ex-60% 2023 and 2024) from the Università degli Studi di Torino. DD was supported by an AvH-Stiftung awarded by the Alexander von Humboldt Foundation.

